# Distinctive Whole-brain Cell Types Predict Tissue Damage Patterns in Thirteen Neurodegenerative Conditions

**DOI:** 10.1101/2023.06.08.544227

**Authors:** Veronika Pak, Quadri Adewale, Danilo Bzdok, Mahsa Dadar, Yashar Zeighami, Yasser Iturria-Medina

## Abstract

For over a century, brain research narrative has mainly centered on neuron cells. Accordingly, most neurodegenerative studies focus on neuronal dysfunction and their selective vulnerability, while we lack comprehensive analyses of other major cell types’ contribution. By unifying spatial gene expression, structural MRI, and cell deconvolution, here we describe how the human brain distribution of canonical cell types extensively predicts tissue damage in thirteen neurodegenerative conditions, including early-and late-onset Alzheimer’s disease, Parkinson’s disease, dementia with Lewy bodies, amyotrophic lateral sclerosis, mutations in presenilin-1, and three clinical variants of frontotemporal lobar degeneration (behavioural variant, semantic and non-fluent primary progressive aphasia) along with associated 3-repeat and 4-repeat tauopathies and TDP43 proteinopathies types A and C. We reconstructed comprehensive whole-brain reference maps of cellular abundance for six major cell types and identified characteristic axes of spatial overlapping with atrophy. Our results support the strong mediating role of non-neuronal cells, primarily microglia and astrocytes, in spatial vulnerability to tissue loss in neurodegeneration, with distinct and shared across-disorders pathomechanisms. These observations provide critical insights into the multicellular pathophysiology underlying spatiotemporal advance in neurodegeneration. Notably, they also emphasize the need to exceed the current neuro-centric view of brain diseases, supporting the imperative for cell-specific therapeutic targets in neurodegeneration.

## Introduction

Neurodegenerative diseases are characterised by substantial neuronal loss in both the central and peripheral nervous systems^1^. In dementia-related conditions like Alzheimer’s disease (AD), frontotemporal dementia (FTD), and dementia with Lewy bodies (DLB), neurodegeneration can lead to progressive damage in brain regions related with memory, behaviour, and cognition^2^. Other diseases are thought to primarily affect the locomotor system, including motor neurons in amyotrophic lateral sclerosis (ALS) and nigrostriatal dopaminergic circuitry in Parkinson’s disease (PD)^3^. Although each disorder has its own distinct etiology, progression, affected brain areas, and clinical manifestations, recent studies support that most of them share same molecular and cellular mechanisms^4–7^.

While research has been mainly focused on neuronal dysfunction, other brain cells such as astrocytes, microglia, oligodendrocytes, as well as cells of the vascular and peripheral immune systems, are gaining more recognition for their contribution to disease pathology^8–10^. Depending on the disease stage, non-neuronal cells in the brain can play a dual role, with their complex response having both protective and detrimental effects on neuronal health and survival^11, 12^. For instance, such glial cells as astrocytes and microglia are involved in neuronal support, maintenance of extracellular homeostasis, and immune regulation in response to injury^13, 14^. Initially, these cells respond to injury by releasing neuroprotective neurotrophic factors and antioxidants^13, 14^. However, under certain conditions, prolonged microglial activation can induce reactive astrocytes and together they release neurotoxic pro-inflammatory cytokines and chemokines, which in turn can lead to metabolic stress and foster the accumulation of amyloid-β and tau plaques in AD, ultimately contributing to heightened neuronal death^15–17^. Growing evidence suggests that immune and other cell type-mediated events are a driving force behind the wide range of neurodegenerative conditions^15, 18–21^. Yet, the exact bases behind how these processes contribute to selective neuronal loss across brain regions remain unclear.

Recent studies have suggested that brain spatial patterns in gene expression are associated with regional vulnerability to some neurodegenerative disorders and their corresponding tissue atrophy distributions^22–26^. Comparison of transcriptomic patterns in middle temporal gyrus across various brain diseases showed cell type expression signature unique for neurodegenerative diseases^7^. Although single-cell transcriptomics and multiomics analyses have advanced our knowledge of cell type compositions associated with pathology in neurodegeneration^27–29^, these are invariably restricted to a few isolated brain regions, usually needing to be preselected at hand for each specific disease. Due to the invasive nature of tissue acquisition/mapping and further technical limitations for covering extended areas^30^, no whole-brain maps for the abundance of cell populations in humans are currently available, constraining the analysis of large-scale cellular vulnerabilities in neurological diseases. Accordingly, how spatial cell types distributions relate to stereotypic regional damages in different neurodegenerative conditions remain largely unclear^31^.

Here, we extend previous analyses of cellular-based spatiotemporal vulnerability in neurodegeneration in three fundamental ways. First, we use transcriptomics, structural magnetic resonance imaging (MRI), and advanced cell deconvolution to construct whole-brain reference maps of cellular abundance in healthy humans for six canonical cell types: neurons, astrocytes, oligodendrocytes, microglia, endothelial cells, and oligodendrocyte precursors. Second, we describe the spatial associations of each healthy level of reference canonical cell types with atrophy in thirteen low-to-high prevalent neurodegenerative conditions, including early-and late-onset AD, genetic mutations in presenilin-1 (PS1 or PSEN1), DLB, ALS, PD, and both clinical and pathological subtypes of frontotemporal lobar degeneration (FTLD). Third, we identify distinctive cell-cell and disorder-disorder axes of spatial susceptibility in neurodegeneration, obtaining new insights about across-disorders (dis)similarities in underlying pathological cellular systems. We confirm that non-neuronal cells express substantial vulnerability to tissue loss and spatial brain alterations in most studied neurodegenerative conditions, with distinct and shared across-cells and across-disorders mechanisms. This study aids in unraveling the commonalities across a myriad of dissimilar neurological conditions, while also revealing cell type specific patterns conferring increased vulnerability or resilience to each examined disorder. For further translation and validation of our findings, all resulting analytic tools and cells abundance maps are shared with the scientific and clinical communities.

## Results

### Multimodal data origin and unification approach

We obtained whole-brain voxel-wise atrophy maps for thirteen neurodegenerative conditions, including early-and late-onset Alzheimer’s disease (EOAD and LOAD, respectively), Parkinson’s disease (PD), amyotrophic lateral sclerosis (ALS), dementia with Lewy bodies (DLB), mutations in presenilin-1 (PS-1), clinical variants of frontotemporal dementia (the behavioural variant bvFTD and the non-fluent and semantic variants of primary progressive aphasia nfvPPA and svPPA), and FTLD-related pathologies such as FLTD-TDP (TAR DNA-binding protein) types A and C, 3-repeat tauopathy, and 4-repeat tauopathy (see *Materials and Methods*, *Disease-specific atrophy maps* subsection)^32–35^. We use the term FTD when addressing the clinical syndromes, and the term FTLD is employed when referencing histologically confirmed neurodegenerative pathologies^36^. Pathological diagnosis confirmation was performed for early-and late-onset AD, DLB, PS-1, FTLD-TDP types A and C, 3-repeat tauopathy, and 4-repeat tauopathy^32^, while PD, ALS, and variants of FTD were diagnosed based on clinical and/or neuroimaging criteria^37–39^, with some ALS patients being histologically confirmed *post-mortem*^38^. Changes in tissue density in the atrophy maps were previously measured by voxel-and deformation-based morphometry (VBM and DBM; *Materials and Methods*, *Disease-specific atrophy maps* subsection) applied to structural T1-weighted MR images, and expressed as a t-score per voxel (relatively low negative values indicate greater GM tissue loss/atrophy; ^40, 41^). All maps are registered to the Montreal Neurological Institute (MNI) brain space^42^. In addition, we obtained bulk transcriptomic data for the adult healthy human brains from the Allen Human Brain Atlas (AHBA)^43^. This included high-resolution coverage of nearly the entire brain, measuring expression levels for over 20,000 genes from 3702 distinct tissue samples of six post-mortem specimens, and detailed structural MRI data (see *Materials and Methods*, *Mapping gene expression data*)^43^.

Using a previously validated approach to infer gene expression levels (in AHBA data) at not-sampled brain locations with Gaussian process regression^44^, mRNA expression levels were completed for all grey matter (GM) voxels in the standardized MNI brain space^42^. Gaussian process regression allowed predicting gene expression values for unobserved regions based on the mRNA values of proximal regions. Next, at each GM location, densities for multiple canonical cell types were estimated using the Brain Cell-type Specific Gene Expression Analysis software (BRETIGEA)^45^. The deconvolution method^45, 46^ (implemented in the BRETIGEA) accurately estimated cell proportions from bulk gene expression for six major cell types (Fig. 1C): neurons, astrocytes, oligodendrocytes, microglia, endothelial cells, and oligodendrocyte precursor cells (OPCs). Overall, atrophy levels for thirteen neurodegenerative conditions and proportion values for six major cell types from healthy brains were unified at matched and standardized locations (MNI space), covering the entire grey matter of the brain (see Fig. 1 for schematic description).

**Figure 1.**
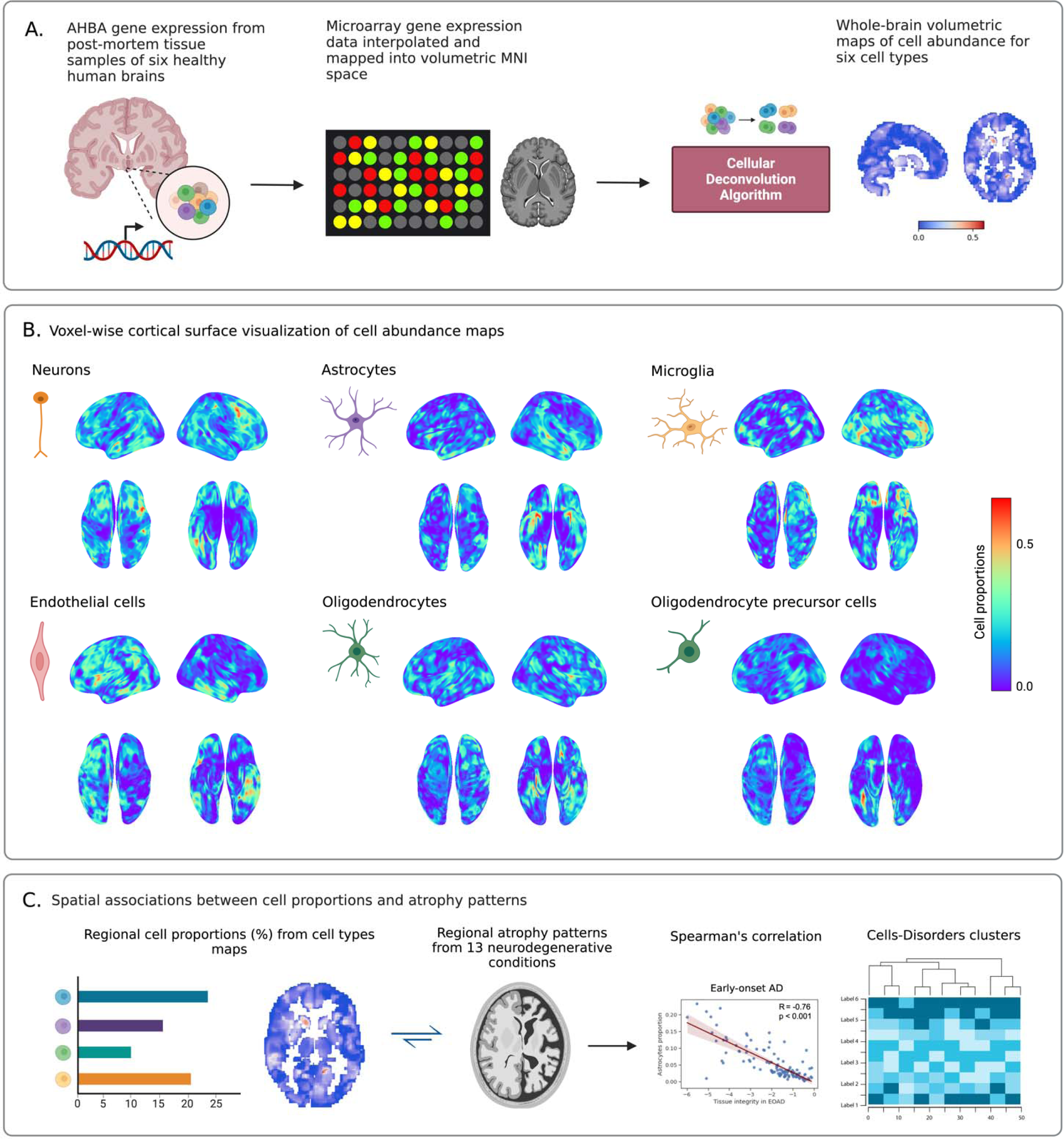
Schematic approach for whole-brain cell type proportions vulnerability analysis in neurodegeneration. (**A**) Microarray bulk gene expression levels in the AHBA were derived from 3072 distinct tissue samples of six post-mortem healthy human brains. Missing gene expression data were then inferred for each unsampled grey matter voxel using Gaussian process regression. When combined with original AHBA data, they were mapped into volumetric MNI space, resulting in the whole-brain transcriptional atlas. Deconvolution algorithm for bulk RNA expression levels was applied to the transcriptional atlas with using well-known cell type-specific gene markers to estimate cell type proportions. Comprehensive volumetric maps showing reconstructed distributions of six canonical cell types across all grey matter voxels in the brain were created (see *Materials and Methods*, *Cell Type proportion estimation* subsection). (**B**) Voxel-wise surface visualization (lateral, dorsal, and ventral views) of cell abundance maps for neurons, astrocytes, microglia, endothelial cells, oligodendrocytes, and OPCs. At each voxel, red and blue colors indicate high and low proportion densities, respectively. (**C**) Associations between cell type proportions from each density map and atrophy values in thirteen neurodegenerative conditions were analysed in 118 grey matter regions predefined by the AAL atlas. Diagram was created with BioRender.com.

We hypothesized (and tested in next subsections) that brain tissue damages in neurodegenerative conditions are associated with distinctive patterns of cells distributions, with alterations on major cell types playing a key role on the development of each disorder and representing a direct factor contributing to brain dysfunction.

### Uncovering spatial associations between cell type abundances and tissue damage in neurodegeneration

First, we investigated whether stereotypic brain atrophy patterns in neurodegenerative conditions show systematic associations with the spatial distribution of canonical cell type populations in healthy brains. For each condition and cell type pair, the non-linear Spearman’s correlation coefficient was calculated with paired atrophy-cell proportion values across 118 cortical and subcortical regions defined by the automated anatomical labelling (AAL) atlas (Table S1; ^47^). The results (Figs. 2A-M and 3A) show clear associations for all the studied conditions, suggesting extensive cell types related tissue damage vulnerability in neurodegenerative conditions. We confirmed that the observed relationships are independent of brain parcellation, obtaining equivalent results for a different brain parcellation (i.e., DKT atlas ^48^; see Fig. S1).

**Figure 2.**
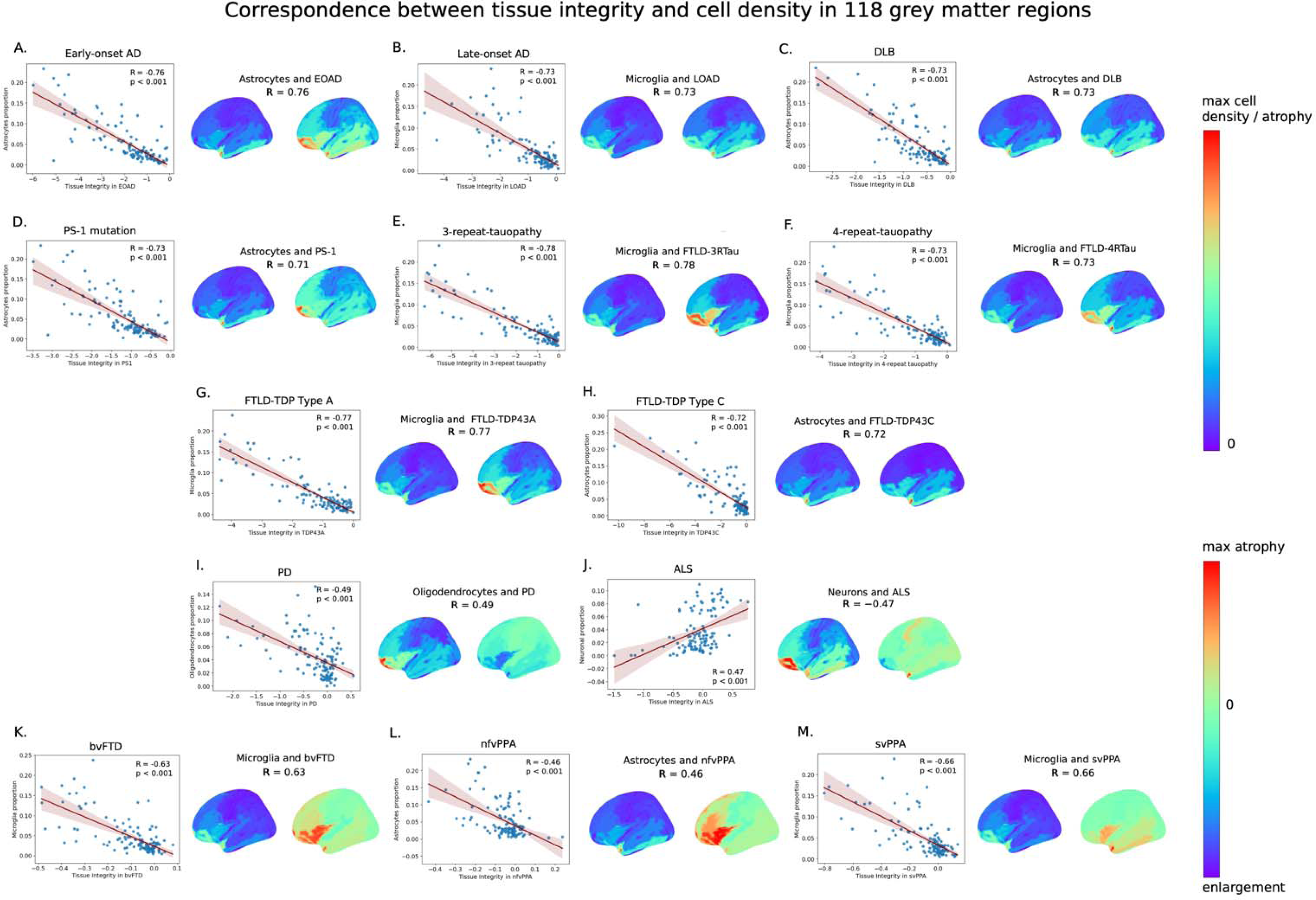
Spatial associations between tissue integrity and cell type proportions for thirteen neurodegenerative conditions illustrated in the scatterplots and surface maps (left hemisphere; lateral view) of regional measures. (**A-M**) Strongest Spearman’s correlations for EOAD, LOAD, DLB, PS1, FTLD-3Rtau, FTLD-4Rtau, FTLD-TDP43A, FTLD-TDP43C, PD, ALS, bvFTD, nfvPPA, and svPPA, respectively. Atrophy and cell type density measures were averaged across 118 grey matter regions and projected to the cortical surface of the *fsaverage* template. Each dot in the scatterplots represents a GM region from the AAL atlas (Table S1; see Fig. S1 for equivalent results for the DKT parcellation). Lower tissue integrity score in the scatterplots’ x-axis indicates greater GM loss/atrophy. For a better visual comparison of patterns in atrophy and cell abundance, the atrophy scale was reversed, with higher t-statistic values indicating greater atrophy in the surface plots. Thus, the first color bar ranging from 0 is universal for all cell maps and pathologically confirmed dementia conditions (A-H). Second color bar captures the tissue enlargement in PD, ALS, and variants of FTD (I-M). Notice how astrocyte density significantly correlates with increase in tissue loss in EOAD, DLB, PS1, FTLD-TDP43C, and nfvPPA (A, C, D, H, L; p < 0.001). Tissue loss was also associated with increase in microglial proportion in LOAD, FTLD-3Rtau, FTLD-4Rtau, FTLD-TDP43A, bvFTD, and svPPA (B, E, F, G, K, M; p < 0.001). Increased oligodendrocytes associated with PD (I; p < 0.001). Increase in neuronal proportion showed association with decrease in atrophy and tissue enrichment in ALS (J; p < 0.001). All p-values were FDR-adjusted with the Benjamini-Hochberg procedure (p < 0.05).

As shown in Figs. 2A-M and 3A, astrocytes and microglia cell occurrences presented the strongest spatial associations with atrophy in most neurodegenerative conditions, particularly for EOAD, LOAD, DLB, PS1, FTLD-3RTau, FTLD-4Rtau, FTLD-TDP type A, FTLD-TDP type C, bvFTD, nfvPPA, and svPPA (all p < 0.001, FDR-corrected). Astrocytes are involved in neuronal support, extracellular homeostasis, and inflammatory regulation in response to injury, and show high susceptibility to senescence and oxidative damage^49, 50^. Astrocytes also play an important role in the maintenance of the blood-brain barrier (BBB), which regulates the passage of molecules, ions, and cells between the blood and the brain^51^. Recent study suggests reactive astrocytes may promote vascular inflammation in the BBB^52^. Endothelial cells, which comprise the functional component of the BBB, also showed strong spatial associations with atrophy in almost all conditions (Fig. 3A). Endothelial cells regulate cerebral blood flow and deliver oxygen and nutrients to the brain^53^. Disruption of the BBB may allow harmful substances to enter the brain, including inflammatory molecules and toxic aggregated proteins, ultimately exacerbating neuronal damage^54, 55^. Reduction of cerebral blood flow and vascular dysregulation are the earliest and strongest pathologic biomarkers of LOAD, PD and other neurodegenerative disorders^56–58^.

**Figure 3.**
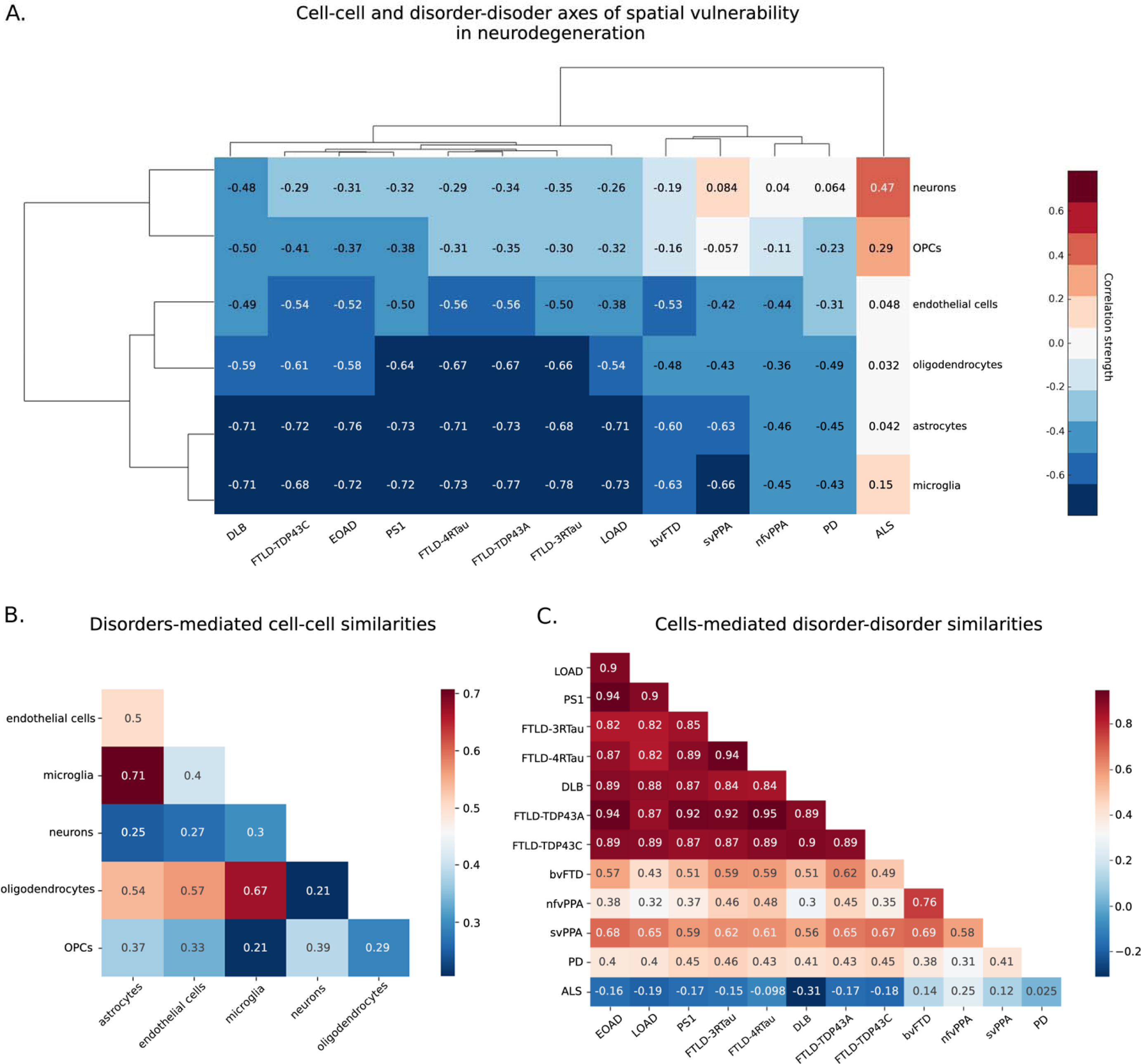
Cells and disorders similarities based on shared distributions. (**A**) Dendrogram and unsupervised hierarchical clustering heatmap of Spearman’s correlations between cell type proportions and atrophy patterns across the thirteen neurodegenerative conditions. (**B**) Cell-cell associations based on regional vulnerabilities to tissue loss across neurodegenerative conditions. (**C**) Disorder-disorder similarities across cell types. In A), red color corresponds to strong positive correlations between cells and disorders, white to no correlation, and dark blue to strong negative correlations.

Similar to astrocytes in their role of supporting neurons, microglial cells are the resident macrophages of the central nervous system and key players in the pathology of neurodegenerative conditions, including AD, PD, FTD and ALS^11, 59, 60^. Besides its many critical specializations, microglial activation in prolonged neuroinflammation is of particular relevance in neurodegeneration^11,61^. At earlier stages of AD, increased population of microglia and astrocytes (microgliosis and astrogliosis) have been observed in diseased regions, due to sustained cellular proliferation in response to disturbances, loss of homeostasis or the accumulation of misfolded proteins^14, 62, 63^. Excessive proliferation may lead to the transition of homeostatic microglia to its senescent or disease-associated type, also known as DAM, via the processes mediated by *TREM2-APOE* signalling^62, 64, 65^. Increased number of dystrophic microglia, a form of cellular senescence characterized as beading and fragmentation of the branches of microglia, has been seen in multiple neurodegenerative conditions such as AD, DLB and TDP-43 encephalopathy^66^. The presence of senescent microglia is believed to ultimately contribute to the failure of brain homeostasis and to clinical symptomatology^21, 65, 67^.

Oligodendrocytes also associated with spatial tissue vulnerability to all conditions aside ALS (Fig. 3A). Oligodendrocytes are responsible for the synthesis and maintenance of myelin in the brain^68^. Demyelination produces loss of axonal insulation leading to neuronal dysfunctions^68, 69^. Myelin dysfunction may lead to secondary inflammation and subsequent failure of microglia to clear amyloid-β deposition in AD mice models^70^. Oligodendrocytes were shown to be highly genetically associated with PD^71–73^. In addition, densities of OPCs showed strong correlations with the atrophy patterns of DLB, EOAD, PS1, and FTLD-TDP type C. OPCs regulate neural activity and harbor immune-related and vascular-related functions^74^. In response to oligodendrocyte damage, OPCs initiate their proliferation and differentiation for the purpose of repairing damaged myelin^75^. In AD, PD and ALS, the OPCs become unable to differentiate and their numbers decrease, leading to a reduction in myelin production and subsequent neural damage^76, 77^.

We observed (Fig. 3A) that neuronal abundance distribution is also associated with tissue damage in many neurodegenerative conditions. However, these associations are less strong than for other cell types, except for the ALS case (Fig. 2J). For this disorder, neuron proportions positively correlated with tissue integrity (i.e., the higher the neuronal proportion, the less atrophy in a region). This observation suggests that increased neuronal presence at brain regions (relative to all considered cell types) may have a protective effect in ALS, making neuronal enriched regions less vulnerable to damage in this disorder. In addition, we observed particularly weak associations between neuronal proportions and tissue damage in all three clinical variants of FTD (bvFTD, nfvPPA, svPPA) and PD (Fig. 3A), suggesting that these conditions may be primary associated with supportive cell types (microglia, astrocytes, and oligodendrocytes, respectively; Figs. 2I, K-M).

### Spatial cell types grouping exposes distinctive disease-disease similarities

Next, we hypothesized that disorders sharing similar biological mechanisms and clinical manifestations present common across-brain patterns of cell type density associations. Fig. 3A shows a hierarchical taxonomy dendrogram grouping cell types and conditions according to their common brain-wide correlation patterns.

The clustergram analysis revealed distinct grouping patterns among various neurodegenerative conditions. All histologically confirmed dementia conditions formed a separate cluster. Notably, EOAD and mutations in PS1, a prevalent cause of familial early-onset AD^78^, grouped together. Interestingly, three clinical subtypes of FTD (bvFTD, nfvPPA, and svPPA) displayed similar patterns of cell type vulnerabilities and diverged into a discrete cluster with PD, separately from ALS and other dementia conditions. However, FTLD-associated pathologies such as TDP-43 proteinopathies (types A and C), as well as 3-repeat and 4-repeat tauopathies, showed patterns more similar to those found in DLB and AD-related conditions (EOAD, LOAD, PS-1) rather than clinical FTD subtypes from a different dataset. These differences could be attributed to variations in the source dataset; the atrophy maps were derived from different studies and were measured by different techniques, which may have introduced discrepancies in results due to different data acquisition tools and protocols (see Methods). Nonetheless, all FTLD-related subtypes and conditions showed strongest associations with spatial distributions of glial cells, particularly astrocytes and microglia.

Among all cell types, neurons and OPCs spatial density distributions were least associated with tissue atrophy in all thirteen conditions, subsequently clustering together. Astrocytes and microglia distributions similarly showed the strongest associations with all neurodegenerative conditions (Fig. 3B), and thus formed a separate cluster while still being related with oligodendrocytes and endothelial cells. Astrocytes and microglia are known to be intimately related in the pathophysiological processes of neurodegenerative disorders^14^. Both are key regulators of inflammatory responses in the central nervous system, and given their role in clearing misfolded proteins, dysfunctions of each of them can result in the accumulation of Aβ and tau^14, 79^. During the progression of AD and PD, microglia’s activation can result in an increased capacity to convert resting astrocytes to reactive astrocytes^17^.

Patterns in cellular vulnerability in DLB did not strongly resemble PD without dementia (Fig. 3C), although both conditions involve alpha-synuclein aggregates^80^. Similar observation can be made for ALS and FTLD. Despite the common presence of TDP-43 abnormal accumulations and their strong genetical overlap^81^, ALS did not group together with FTD variants and FTLD-associated pathologies (FTLD-TDP type A, FTLD-TDP type C) based on patterns of cells-atrophy associations. All these conditions are known to be pathologically linked, often arising from either tau or TDP-43 accumulation; for instance, TDP-43 is the usual cause of svPPA and approximately half of bvFTD cases, while the other half of bvFTD patients and many nfvPPA cases are associated with tau pathology^82^. These results emphasize the fundamental role of network topology and other factors beyond the presence of toxic misfolded proteins in developing characteristic tissue loss and cellular vulnerability in neurodegenerative conditions^34, 83–85^.

## Discussion

Previous efforts to describe the composition of the brain’s different cell populations related to neurodegeneration have been limited to a few isolated regions. In the most systematic study of its kind, here we characterized large-scale spatial associations between canonical cell types and brain tissue loss across cortical and subcortical grey matter areas in thirteen neurodegenerative conditions (including early-and late-onset AD, PD, DLB, ALS, mutations in PS1, and clinical (bvFTD, nfvPPA, svPPA) and pathological (3-repeat and 4-repeat tauopathies and TDPP43 proteinopathies types A and C) subtypes of FTLD. Starting from healthy brain levels of gene expression and structural MRI data from the Allen Human Brain Atlas^43^, and extending our analysis with advanced single cell-RNA seq-validated cell deconvolution approaches, along with whole-brain atrophy maps from clinically and/or neuropathologically confirmed disorders, we determined that (i) the spatial distributions of non-neuronal cell types, primarily microglia and astrocytes, are strongly associated with the spread tissue damage present in many neurodegenerative conditions; (ii) cells and disorders define major axes that underlie spatial vulnerability, aiding in comprehending heterogeneity behind distinct and similar clinical manifestations/definitions; (iii) the generated whole-brain maps of cellular abundance can be similarly used for studying associations between imaging phenotypes and healthy reference cellular levels in other neurological conditions (e.g., neurodevelopmental and neuropsychiatric disorders). Overall, our findings stress the critical need to surpass the current neuro-centric view of brain diseases and the imperative for identifying cell-specific therapeutic targets in neurodegeneration. For further translation and validation, all resulting cells abundance maps and analytic tools are freely shared with the community.

We derived, first to our knowledge, high resolution maps of cellular abundance/proportion in the adult human healthy brain for six canonical cell types, including astrocytes, neurons, oligodendrocytes, microglia, and endothelial cells. As mentioned, previous cellular analyses of neurological conditions have been restricted to expert-selected isolated brain areas. The invasive nature of expression assays, requiring direct access to neural tissue, and other numerous scaling limitations have impeded extensive spatial analyses^86^. Earlier studies, also using AHBA data, have shown that spatial patterns in cell type-specific gene expression are associated with both regional vulnerability to neurodegeneration and patterns of atrophy across the brain^7, 22–25^. Since many neurodegeneration-related genes have similar levels of expression in both affected and unaffected brain areas^87^, characterizing changes in tissue loss associated with reference cell type proportions in health may provide a clearer perspective on large-scale spatial patterns of cellular vulnerability. Our maps of cells-abundance are available for the scientific and clinical community, potentially allowing researchers to further study spatial variations in cell type density with macroscale phenotypes. These maps can be used in future studies concerning brain structure and function in both health and disease. They can be also explored in context of other neurological diseases, including neurodevelopmental and psychiatric conditions.

Our results demonstrate that all canonical cell types express vulnerability to dementia-related atrophy of brain tissue, potentially suggesting the disruption of the molecular pathways involving specific cell types can contribute to their observed dysfunctions and subsequent clinical symptamology^21^. Previously, transcriptional profiling of prefrontal cortex in AD showed reduced proportions of neurons, astrocytes, oligodendrocytes, and homeostatic microglia^88^. In contrast, bulk-RNA analysis of diseased AD tissues from various human brain regions observed neuronal loss and increased cell abundance of microglia, astrocytes, oligodendrocytes, and endothelial cells^89, 90^. Furthermore, increased microglial, endothelial cells, and oligodendrocytes population was observed in PD and other Lewy diseases^72, 91^. Cortical regions exhibiting the most severe atrophy in symptomatic *C9orf72*, *GRN* and *MAPT* mutation carriers with FTD showed increased gene expression of astrocytes and endothelial cells^25^. Cortical thinning has been demonstrated to correlate with higher proportions of astrocytes, microglia, oligodendrocytes, oligodendrocyte precursor cells, and endothelial cells in cases of AD compared to controls^22, 26^. In line with these results, we observed that regions with increased cell type proportions, particularly for astrocytes and microglia, are strongly associated with gray matter atrophy in almost all neurodegenerative conditions. This may partly explain the reported cellular proliferation through microglial activation in diseased regions in response to the misfolded protein accumulation or other pathobiological processes^62, 67^. As disease progresses, the release of inflammatory agents by sustained microglial activation is believed to be responsible for exacerbating neurodegeneration and clinical symptoms^15, 18^. Microglial activation in pair with grey matter atrophy in frontal cortex were shown to be directly associated with cognitive decline in FTD^60^.

Our study has several limitations. Firstly, our analyses were focused on stereotypic atrophy patterns for each disorder. It is known that neurodegenerative diseases are highly heterogenous, with molecular, phenotypic, and clinical subtypes potentially varying in atrophy patterns^92, 93^. Further investigation of cell type signatures across various subtypes not covered in this study and disease stages may better characterize each case. Additionally, comparing our findings with neuropathological assessments of diseased brain tissues in available regions would be beneficial. While the diagnosis of most dementia conditions used in this study has been histologically confirmed, the diagnosis for clinical variants of FTD, ALS, and PD patients was based on clinical and neuroimaging assessments. In addition, it has been observed that cell type-related transcriptional changes are different between sexes^94^, making future sex-specific analyses indispensable for further understanding of sex-related pathomechanisms. An important consideration is that examined atrophy maps were sourced from different studies (Table S2), with differences in data acquisition protocols (e.g., spatial resolution) and technical procedures (e.g., smoothing level, statistical methods). In complementary analyses, we observed almost identical results after smoothing all disorder-specific images with the same kernel size, while they were already mapped at the same spatial resolution for this study and statistically adjusted by acquisition parameters (e.g., field strength) in original studies. Moreover, cell type deconvolution approaches are varied and limited in their precision^95^. Here, we used a previously validated deconvolution method designed for efficiently estimating cell proportions for six major cell types from bulk mRNA expression^45^. Conveniently, this method is freely available for researchers (R package, BRETIGEA), which will facilitate reproducibility analyses of our study. Other important considerations are the dynamic nature of gene expression as disease progresses^96, 97^, *post-mortem* RNA degradation of the used templates^98^, and the subsequent limited ability of bulk RNA sequencing to reflect cell-to-cell variability, which is relevant for understanding cell heterogeneity and the roles of specific cell populations in disease ^99^. Lastly, a promising future direction would be to validate our findings with single-cell spatial analyses.

## Materials and Methods

### Disorder-specific atrophy maps

Voxel-wise brain atrophy maps in early-and late-onset Alzheimer’s disease (EOAD and LOAD), Parkinson’s disease (PD), amyotrophic lateral sclerosis (ALS), dementia with Lewy bodies (DLB), mutations carriers in presenilin-1, clinical variants of frontotemporal dementia (FTD), and frontotemporal lobar degeneration (FTLD) pathologies (FTLD-TDP types A and C, 3-repeat tauopathy, and 4-repeat tauopathy) were adopted from open data repositories and/or requested from collaborators^32–35^, as specified below. Reduction in grey matter (GM) density in diseased atrophy maps relative to controls was measured by voxel-and deformation-based morphometry (VBM and DBM) applied to structural T1-weighted MR images, and thus was expressed as t-score per voxel (relatively low negative t-scores indicate greater GM tissue loss/atrophy)^40, 41^. VBM is a hypothesis-free technique for analyzing neuroimaging data that characterizes regional tissue concentration differences across the whole brain, without the need to predefine regions of interest^100^. DBM is a similar widely used technique to identify structural changes in the brain across participants, which in addition considers anatomical differences such as shape and size of brain structures^101^. See Table S2 for study-origin, sample size and imaging technique corresponding to each atrophy map.

MRI data for neuropathological dementias were collected from 186 individuals with a clinical diagnosis of dementia and histopathological (post-mortem or biopsy) confirmation of underlying pathology, along with 73 healthy controls^32^. Data were averaged across participants per condition: 107 had a primary AD diagnosis (68 early-onset (<65 years at disease onset), 29 late onset (≥65 years at disease onset), 10 presenilin-1 mutation carriers), 25 with DLB, 11 with 3-repeat-tauopathy, 17 with 4-repeat-tauopathy, 12 FTLD-TDP type A, and 14 FTLD-TDP type C)^32^. Imaging data were collected from multiple centres on scanners from three different manufacturers (Philips, GE, Siemens), using a variety of different imaging protocols^32^. Magnetic field strength varied between 1.0 T (n=15 scans), 1.5 T (n=201 scans) and 3 T (n=43 scans)^32^. Pathological examination of brain tissue was conducted between 1997 and 2015 according to the standard histopathological processes and criteria in use at the time of assessment at one of four centres: the Queen Square Brain Bank, London; Kings College Hospital, London; VU Medical Centre, Amsterdam and Institute for Ageing and Health, Newcastle^32^. Atrophy maps were statistically adjusted for age, sex, total intracranial volume, and MRI strength field and site^32^. Ethical approval for this retrospective study was obtained from the National Research Ethics Service Committee London-Southeast^32^.

MRI data for PD, consisted of 3T high-resolution T1-weighted scans, were obtained from the Parkinson’s Progression Markers Initiative (PPMI) database (https://www.ppmi-info.org/)^37^. The PPMI is a multi-center international study with approved protocols by the local institutional review boards at all 24 sites across the US, Europe, and Australia^37^. MRI data were acquired in 16 centers participating in the PPMI project, using scanners from three different manufacturers (GE medical systems, Siemens, and Philips medical systems). 3T high-resolution T1-weighted MRI scans from the initial visit and clinical data used in constructing atrophy maps were collected from 232 participants with PD and 118 age-matched controls^34^. PD subjects (77 females; age 61.2 ± 9.1) were required to be at least 30 years old or older, untreated with PD medications, diagnosed within the last two years, and to exhibit at least two or more PD-related motor symptoms, such as asymmetrical resting tremor, uneven bradykinesia, or a combination of bradykinesia, resting tremor, and rigidity^37^. All individuals underwent dopamine transporter (DAT) imaging to confirm a DAT deficit as a prerequisite for eligibility^37^. No significant effect of age, gender, or site was found^34^.

For ALS, MRI data were collected from 66 patients (24 females; age 57.98 ± 10.84) with both sporadic or familial form of disease from centers of the Canadian ALS Neuroimaging Consortium (http://calsnic.org/, ClinicalTrials.gov NCT02405182), which included 3T MRI sites in University of Alberta, University of Calgary, University of Toronto, and McGill University^33, 38^. Patients were included if they were diagnosed with sporadic or familial ALS, and meet the revised El Escorial research criteria^102^ for possible, laboratory supported, or definite ALS^38^. Patients underwent a neurological exam administered by a trained neurologist at each participating site^38^. All participants gave written informed consent, and the study was approved by the health research ethics boards at each of the participating sites^33^. Participants were excluded if they had a history of other neurological or psychiatric disorders, prior brain injury, or respiratory impairment resulting in an inability to tolerate the MRI protocol^33^. Participants with primary lateral sclerosis, progressive muscular atrophy, or frontotemporal dementia were also excluded from the study^38^. Normative aging as well as sex differences were regressed out from data prior the map construction^33^.

For clinical subtypes of FTD, atrophy maps were obtained from the open-access database^35^. These maps were derived from MRI data from the Frontotemporal Lobar Degeneration Neuroimaging Initiative (FTLDNI AG032306; part of the ALLFTD). As described in separate studies^103, 104^, the data used for constructing these atrophy maps consisted of 136 patients diagnosed with frontotemporal dementia, alongside 133 age-matched control participants. Participants were previously stratified into groups according to their clinical variant of FTD: 70 patients were diagnosed with the behavioural variant, 36 with the semantic primary progressive aphasia, and 30 with the non-fluent primary progressive aphasia^39, 103^. 3T structural images were collected on three following sites: University of California San Francisco, Mayo Clinic, and Massachusetts General Hospital^39^. Patients were referred by physicians or self-referred, and all underwent neurological, neuropsychological, and functional assessment with informant interview^39^. All individuals received their diagnoses during a multidisciplinary consensus conference using established criteria: Neary criteria^105^ or, depending on year of enrolment, the recently published consensus criteria for bvFTD^106^ and PPA^107^. Histological analysis was conducted to assess whether patients might have Alzheimer’s disease pathology since both conditions presents the overlap of clinical symptoms^39^. All subjects provided informed consent and the protocol was approved by the institutional review board at all sites^39^.

### Mapping gene expression data

To construct a comprehensive transcriptome atlas, we used mRNA microarray gene expression data from the Allen Human Brain Atlas (AHBA; https://human.brain-map.org/)^43^. The AHBA included anatomical and histological data collected from six healthy human specimens with no known neurological disease history (one female; age range 24–57 years; mean age 42.5 ± 13.38 years)^43^. Two specimens contained data from the entire brain, whereas the remaining four included data from the left hemisphere only, with 3702 spatially distinct samples in total^43^. The samples were distributed across cortical, subcortical, brainstem, and cerebellar regions in each brain, and the expression levels of more than 20,000 genes were quantified^43^. mRNA data for specific brain locations were accompanied by structural MR data from each individual and were labeled with Talairach native coordinates^108^ and Montreal Neurological Institute (MNI) coordinates^42^, which allowed us to match samples to imaging data.

Following the validated approach in ^44^, missing data points between samples for each MNI coordinate were interpolated using Gaussian-process regression, a widely used method for data interpolation in geostatistics. The regression is performed as a weighted linear combination of missing mRNA, with the weights decreasing from proximal to distal regions. MNI coordinates for predicting mRNA values were taken from the GM regions of the AAL atlas. Spatial covariance between coordinates from the available 3072 AHBA tissue samples and coordinates from the AAL atlas was estimated via the quadratic exponential kernel function. mRNA expression at each MNI coordinate was then predicted by multiplying AHBA gene express values that corresponded to specific probes to kernel covariance matrix divided by the sum of kernels.

### Cell type proportion estimation

Densities for multiple canonical cell types were estimated at the grey matter (GM) by applying an R-package Brain Cell-type Specific Gene Expression Analysis (BRETIGEA), with known genetic markers to the transcriptome atlas^45^. This eigengene decomposition-based deconvolution method was designed for estimating cell proportions in bulk gene expression data for six major cell types: neurons, astrocytes, oligodendrocytes, microglia, endothelial cells, and oligodendrocyte precursor cells^45, 46^. We chose 15 representative gene markers per each cell type (90 in total) from the BRETIGEA human brain marker gene set and then selected those genes that were also present in the AHBA gene expression database with matching gene probes. This resulted in eighty cell type-related gene markers that were used in missing data interpolation and the deconvolution proportion estimation analysis (Table S3). For each voxel, each cell type proportion value was normalized relative to the sum of all six cell types and the sum was scaled relative to the grey matter density. We then registered data into MNI and volumetric space using the ICBM152 template^42^.

For the correlation analysis, cell densities were averaged over 118 anatomical regions in grey matter defined by the extended automated anatomical labelling atlas (AAL; Table S1)^47^. We repeated the correlation analysis for the 98 regions from the Desikan-Killiany-Tourville atlas (DKT; Fig. S1)^48^.

## Data analysis

We constructed a 6LJ×LJ13 correlation matrix by computing inter-regional Spearman’s correlations between spatial distributions of the six canonical cell types and patterns of atrophy in thirteen neurodegenerative conditions. Correction for multiple comparisons using the false discovery rate (FDR) was conducted using the Benjamini-Hochberg method, with a significance threshold of 0.05. Shapiro-Wilk tests were used to examine the normality of data distribution. Hierarchical clustering analyses was applied using in-built MATLAB function for data visualization. Cells and conditions were clustered together based on estimated averaged linkage Euclidian distance between their correlation values.

## Data and materials availability

All data needed to evaluate the conclusions in the paper are present in the paper and/or the Supplementary Materials. The BRETIGEA R package can be downloaded from https://cran.r-project.org/package=BRETIGEA. The Allen Human Brain Atlas data is available at https://human.brain-map.org/static/download. Atrophy maps for pathologically confirmed dementia are available at http://neurovault.org/collections/ADHMHOPN/. Raw demographic and MRI data from PD and ALS patients can be accessed at https://www.ppmi-info.org/ and http://calsnic.org/ (ClinicalTrials.gov NCT02405182), respectively. Atrophy maps for clinical variants of FTD are available at https://zenodo.org/records/10383493. Raw data from the FTLDNI initiative can be downloaded from the Laboratory of Neuroimaging (LONI) Image Data Archive at https://ida.loni.usc.edu. The cells abundance maps from this study are freely shared with the community and can be found at our lab’s GitHub space https://github.com/neuropm-lab/cellmaps.

## Acknowledgments

This project was undertaken thanks in part to the following funding awards to YIM: the Canada Research Chair tier 2, the CIHR Project Grant 2020, the Weston Family Foundation’s Transformational Research in AD 2020, and the New Investigator start-up grant from McGill University’s *Healthy Brains for Healthy Lives Initiative* (HBHL, *Canada First Research Excellence Fund*). First author VP was supported by the Laszlo & Etelka Kollar Fellowship from the Faculty of Medicine and Health Sciences at McGill University and partly supported by the HBHL’s Theme 1 Discovery fund 2022-2025. In addition, we used the computational infrastructure of the McConnell Brain Imaging Center at the Montreal Neurological Institute, supported in part by the *Brain Canada Foundation*, through the *Canada Brain Research Fund*, with the financial support of *Health Canada* and sponsors.

## Competing interests

The authors declare no competing interest.

## Supplementary Information

**Figure S1.**
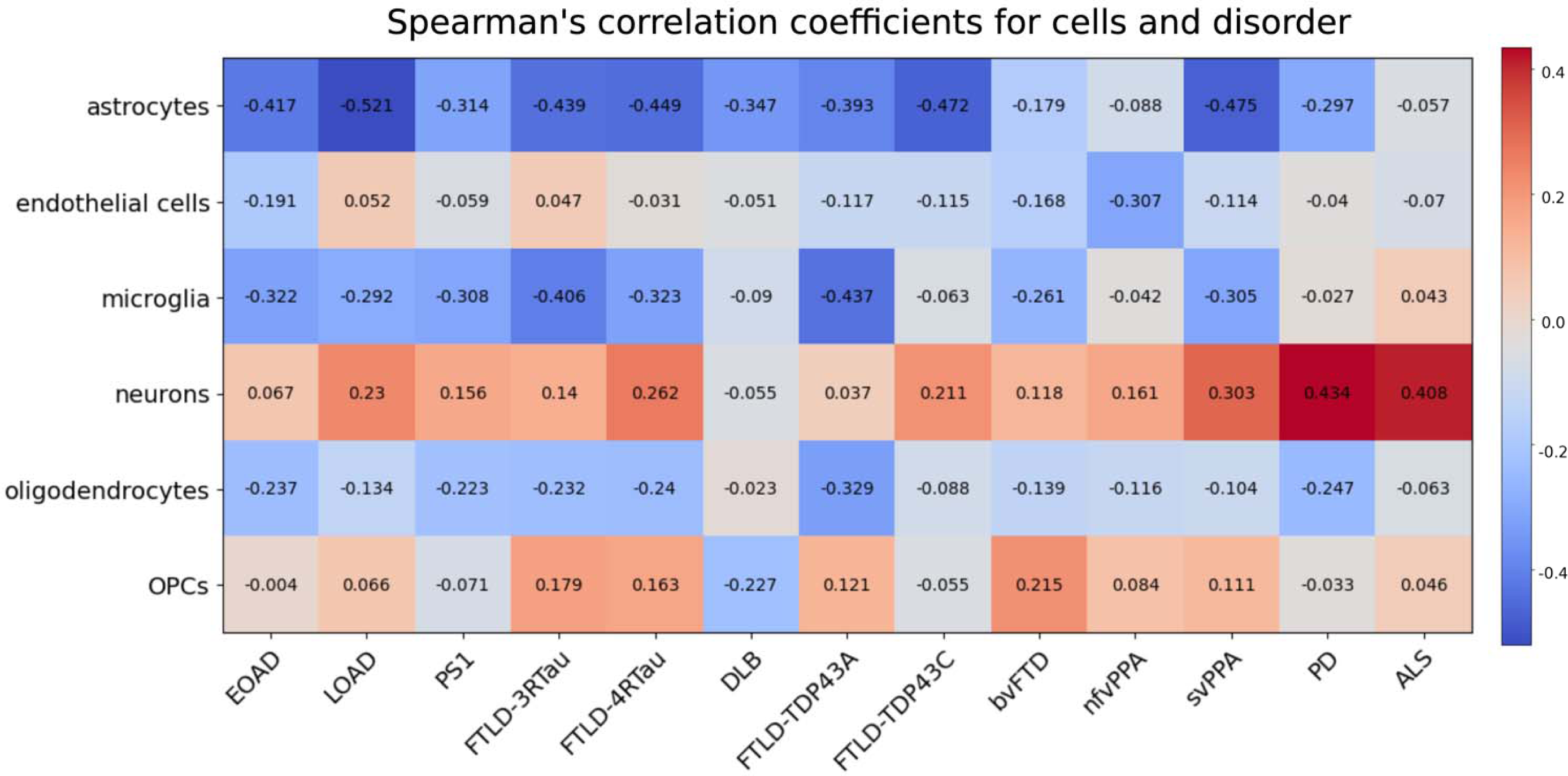
Spatial associations between tissue integrity and cell type proportions for thirteen neurodegenerative conditions in GM regions from the DKT parcellation (equivalent to main results from the AAL atlas in Fig. 3A).

**Table S1.**
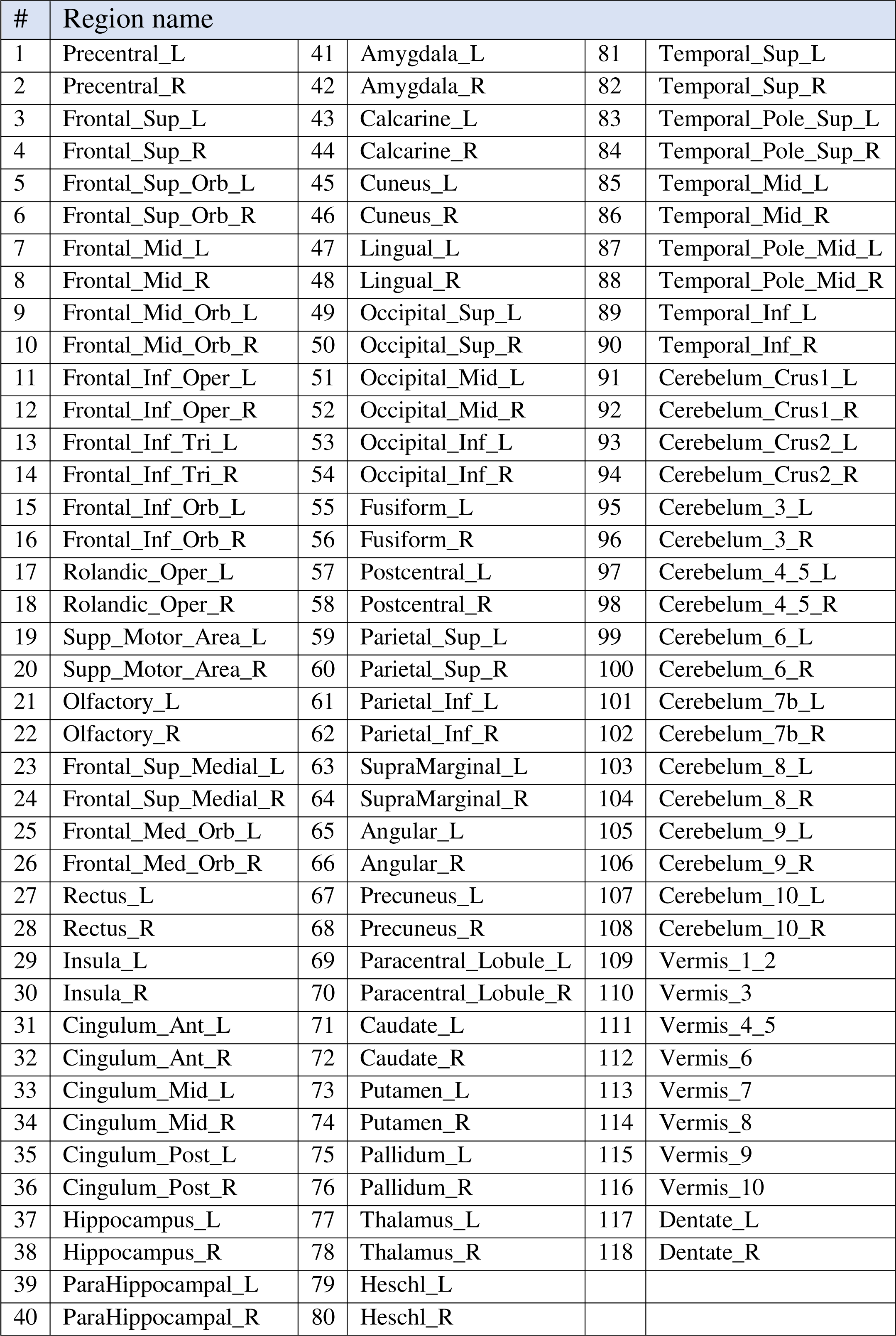
Cortical and subcortical regions from the AAL atlas.

**Table S2.**
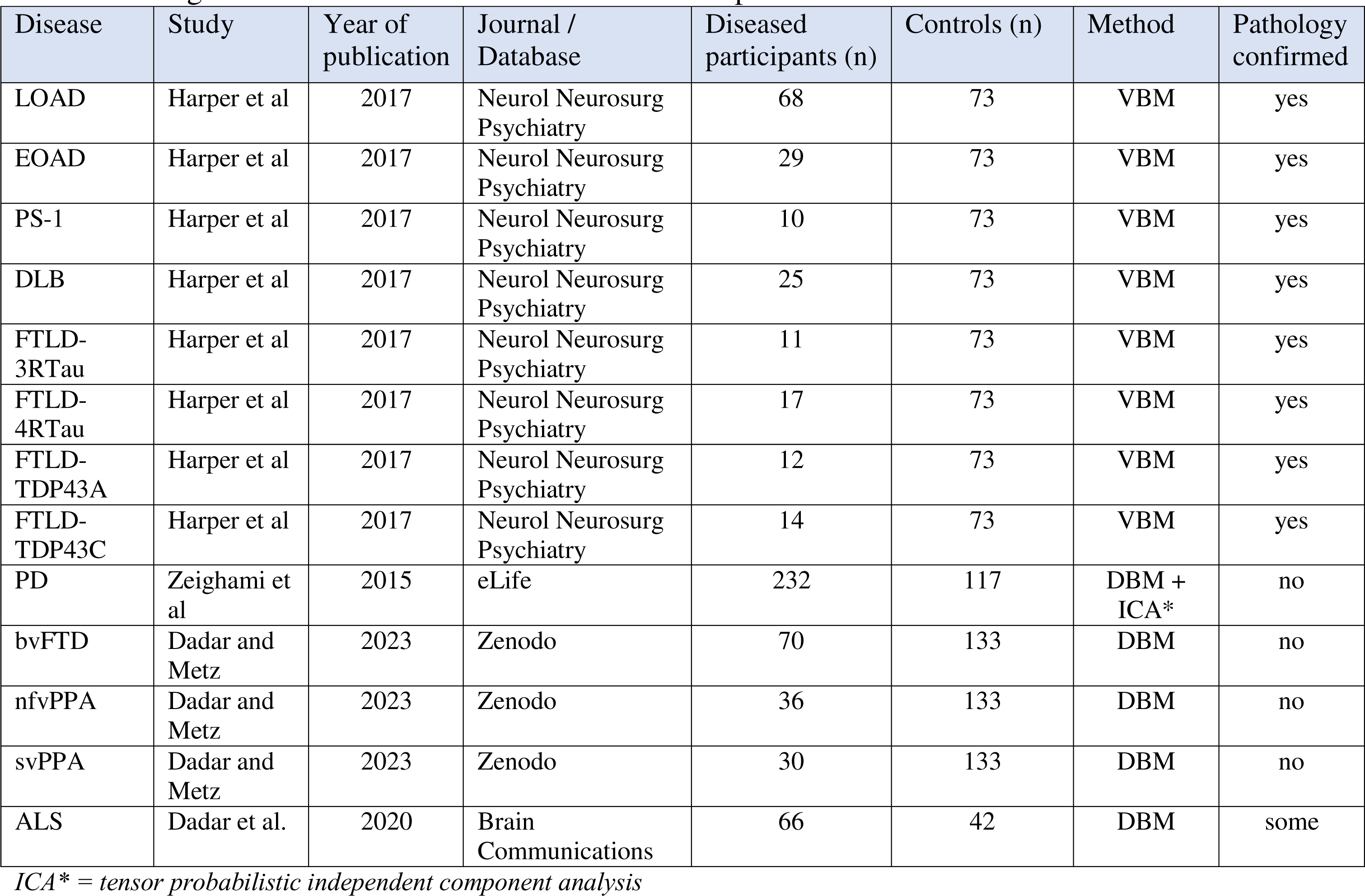
Origin of each disorder-associated t-statistic map.

**Table S3.**
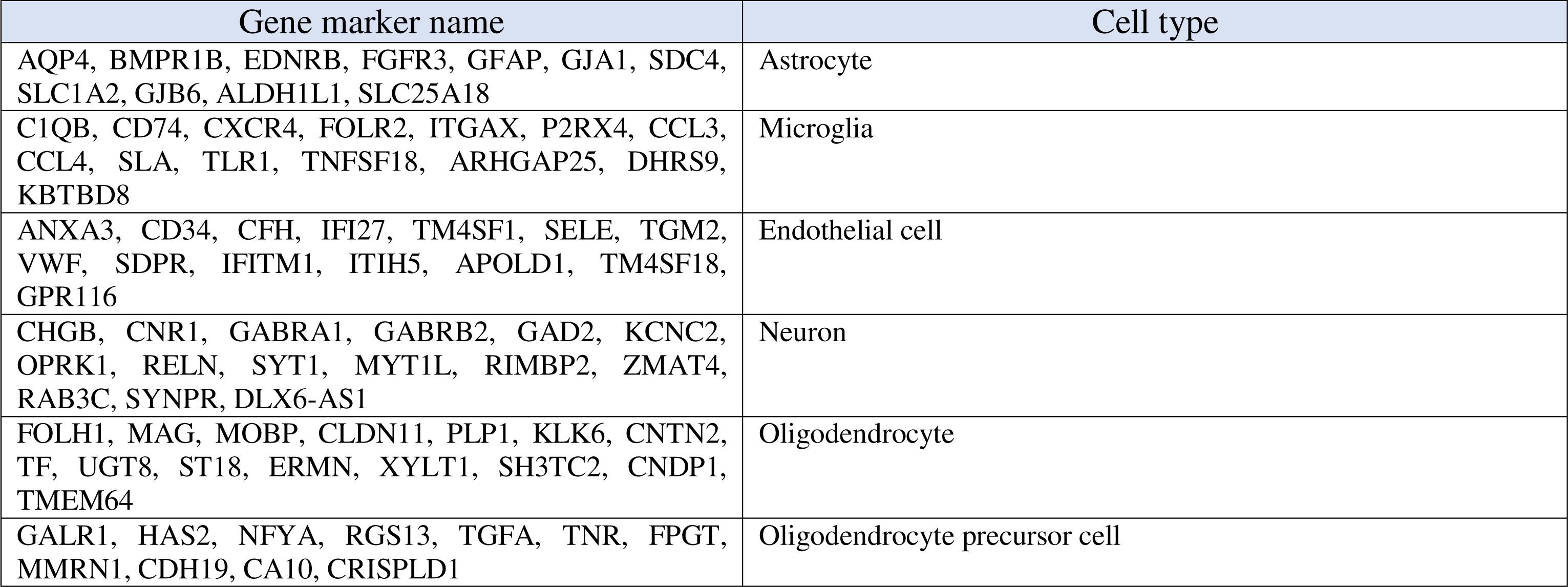
Eighty cell type related gene markers provided by the BRETIGEA R package.

## Notes

### Competing Interest Statement

The authors have declared no competing interest.

### Summary of Updates

The final version of the paper after extensive revision.

