## Supplementary information for "Distinctive Whole-brain Cell Types Predict Tissue Damage Patterns in Thirteen Neurodegenerative Conditions"

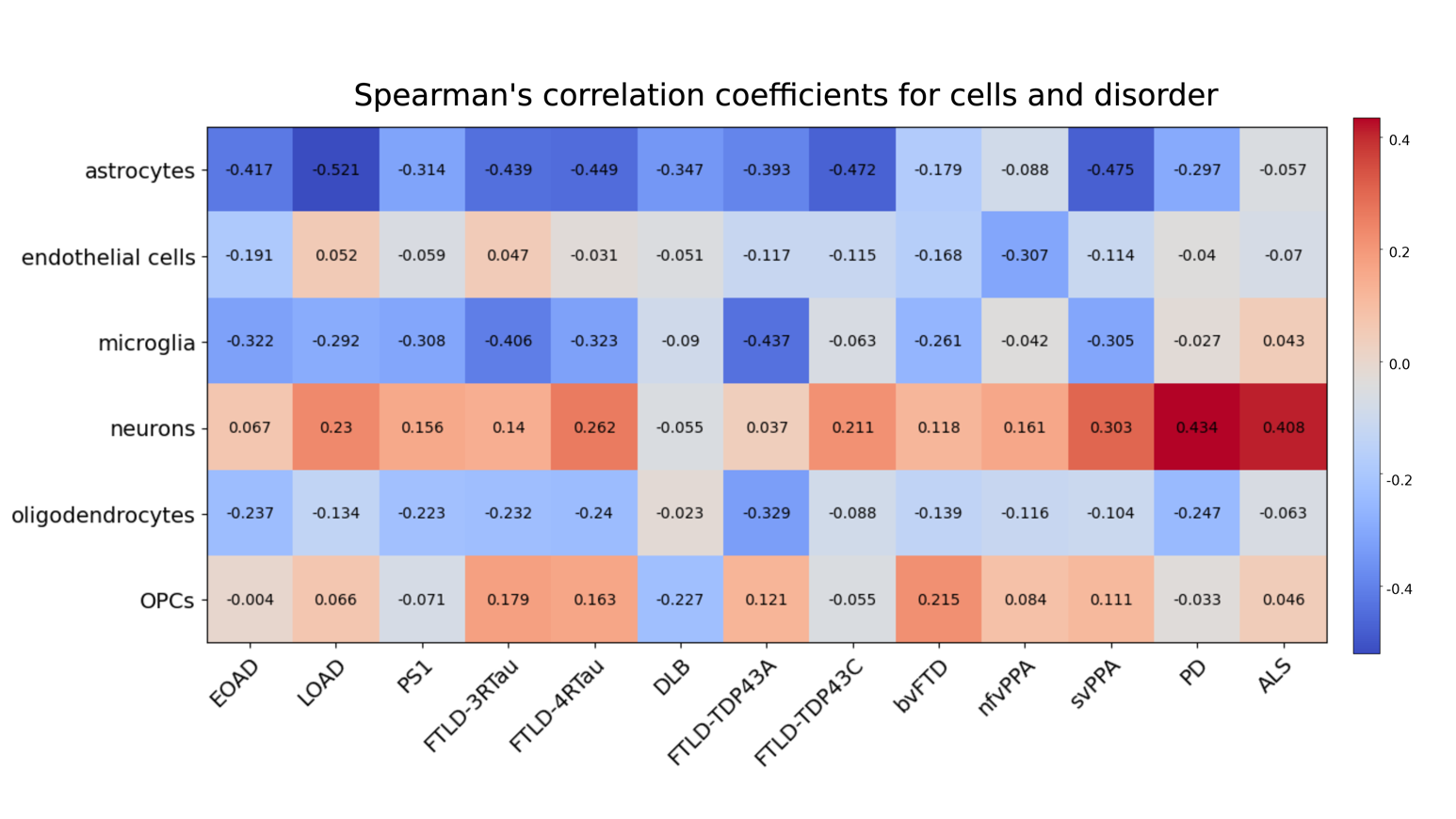

**Figure S1.** Spatial associations between tissue integrity and cell-types proportions for thirteen neurodegenerative conditions in GM regions from the DKT parcellation (equivalent to main results from the AAL atlas in Figure 2).

**Table S1.** Cortical and subcortical regions from the AAL atlas.

| # | Region name | | | | |
| --- | --- | --- | --- | --- | --- |
| 1 | Precentral_L | 41 | Amygdala_L | 81 | Temporal_Sup_L |
| 2 | Precentral_R | 42 | Amygdala_R | 82 | Temporal_Sup_R |
| 3 | Frontal_Sup_L | 43 | Calcarine_L | 83 | Temporal_Pole_Sup_L |
| 4 | Frontal_Sup_R | 44 | Calcarine_R | 84 | Temporal_Pole_Sup_R |
| 5 | Frontal_Sup_Orb_L | 45 | Cuneus_L | 85 | Temporal_Mid_L |
| 6 | Frontal_Sup_Orb_R | 46 | Cuneus_R | 86 | Temporal_Mid_R |
| 7 | Frontal_Mid_L | 47 | Lingual_L | 87 | Temporal_Pole_Mid_L |
| 8 | Frontal_Mid_R | 48 | Lingual_R | 88 | Temporal_Pole_Mid_R |
| 9 | Frontal_Mid_Orb_L | 49 | Occipital_Sup_L | 89 | Temporal_Inf_L |
| 10 | Frontal_Mid_Orb_R | 50 | Occipital_Sup_R | 90 | Temporal_Inf_R |
| 11 | Frontal_Inf_Oper_L | 51 | Occipital_Mid_L | 91 | Cerebelum_Crus1_L |
| 12 | Frontal_Inf_Oper_R | 52 | Occipital_Mid_R | 92 | Cerebelum_Crus1_R |
| 13 | Frontal_Inf_Tri_L | 53 | Occipital_Inf_L | 93 | Cerebelum_Crus2_L |
| 14 | Frontal_Inf_Tri_R | 54 | Occipital_Inf_R | 94 | Cerebelum_Crus2_R |
| 15 | Frontal_Inf_Orb_L | 55 | Fusiform_L | 95 | Cerebelum_3_L |
| 16 | Frontal_Inf_Orb_R | 56 | Fusiform_R | 96 | Cerebelum_3_R |
| 17 | Rolandic_Oper_L | 57 | Postcentral_L | 97 | Cerebelum_4_5_L |
| 18 | Rolandic_Oper_R | 58 | Postcentral_R | 98 | Cerebelum_4_5_R |
| 19 | Supp_Motor_Area_L | 59 | Parietal_Sup_L | 99 | Cerebelum_6_L |
| 20 | Supp_Motor_Area_R | 60 | Parietal_Sup_R | 100 | Cerebelum_6_R |
| 21 | Olfactory_L | 61 | Parietal_Inf_L | 101 | Cerebelum_7b_L |
| 22 | Olfactory_R | 62 | Parietal_Inf_R | 102 | Cerebelum_7b_R |
| 23 | Frontal_Sup_Medial_L | 63 | SupraMarginal_L | 103 | Cerebelum_8_L |
| 24 | Frontal_Sup_Medial_R | 64 | SupraMarginal_R | 104 | Cerebelum_8_R |
| 25 | Frontal_Med_Orb_L | 65 | Angular_L | 105 | Cerebelum_9_L |
| 26 | Frontal_Med_Orb_R | 66 | Angular_R | 106 | Cerebelum_9_R |
| 27 | Rectus_L | 67 | Precuneus_L | 107 | Cerebelum_10_L |
| 28 | Rectus_R | 68 | Precuneus_R | 108 | Cerebelum_10_R |
| 29 | Insula_L | 69 | Paracentral_Lobule_L | 109 | Vermis_1_2 |
| 30 | Insula_R | 70 | Paracentral_Lobule_R | 110 | Vermis_3 |
| 31 | Cingulum_Ant_L | 71 | Caudate_L | 111 | Vermis_4_5 |
| 32 | Cingulum_Ant_R | 72 | Caudate_R | 112 | Vermis_6 |
| 33 | Cingulum_Mid_L | 73 | Putamen_L | 113 | Vermis_7 |
| 34 | Cingulum_Mid_R | 74 | Putamen_R | 114 | Vermis_8 |
| 35 | Cingulum_Post_L | 75 | Pallidum_L | 115 | Vermis_9 |
| 36 | Cingulum_Post_R | 76 | Pallidum_R | 116 | Vermis_10 |
| 37 | Hippocampus_L | 77 | Thalamus_L | 117 | Dentate_L |
| 38 | Hippocampus_R | 78 | Thalamus_R | 118 | Dentate_R |
| 39 | ParaHippocampal_L | 79 | Heschl_L |  |  |
| 40 | ParaHippocampal_R | 80 | Heschl_R |  |  |

**Table S2.** Origin of each disorder-associated t-statistic map.

| Disease | Study | Year of publication | Journal / Database | Diseased participants (n) | Controls (n) | Method | Pathology confirmed |
| --- | --- | --- | --- | --- | --- | --- | --- |
| LOAD | Harper et al | 2017 | Neurol Neurosurg Psychiatry | 68 | 73 | VBM | yes |
| EOAD | Harper et al | 2017 | Neurol Neurosurg Psychiatry | 29 | 73 | VBM | yes |
| PS-1 | Harper et al | 2017 | Neurol Neurosurg Psychiatry | 10 | 73 | VBM | yes |
| DLB | Harper et al | 2017 | Neurol Neurosurg Psychiatry | 25 | 73 | VBM | yes |
| FTLD-3RTau | Harper et al | 2017 | Neurol Neurosurg Psychiatry | 11 | 73 | VBM | yes |
| FTLD-4RTau | Harper et al | 2017 | Neurol Neurosurg Psychiatry | 17 | 73 | VBM | yes |
| FTLD-TDP43A | Harper et al | 2017 | Neurol Neurosurg Psychiatry | 12 | 73 | VBM | yes |
| FTLD-TDP43C | Harper et al | 2017 | Neurol Neurosurg Psychiatry | 14 | 73 | VBM | yes |
| PD | Zeighami et al | 2015 | eLife | 232 | 117 | DBM + ICA* | no |
| bvFTD | Dadar and Metz | 2023 | Zenodo | 70 | 133 | DBM | no |
| nfvPPA | Dadar and Metz | 2023 | Zenodo | 36 | 133 | DBM | no |
| svPPA | Dadar and Metz | 2023 | Zenodo | 30 | 133 | DBM | no |
| ALS | Dadar et al. | 2020 | Brain Communications | 66 | 42 | DBM | some |

*ICA* = tensor probabilistic independent component analysis*

**Table S3.** Eighty cell-type related gene markers provided by the BRETIGEA R package.

| Gene marker name | Cell-type |
| --- | --- |
| AQP4, BMPR1B, EDNRB, FGFR3, GFAP, GJA1, SDC4, SLC1A2, GJB6, ALDH1L1, SLC25A18 | Astrocyte |
| C1QB, CD74, CXCR4, FOLR2, ITGAX, P2RX4, CCL3, CCL4, SLA, TLR1, TNFSF18, ARHGAP25, DHRS9, KBTBD8 | Microglia |
| ANXA3, CD34, CFH, IFI27, TM4SF1, SELE, TGM2, VWF, SDPR, IFITM1, ITIH5, APOLD1, TM4SF18, GPR116 | Endothelial cell |
| CHGB, CNR1, GABRA1, GABRB2, GAD2, KCNC2, OPRK1, RELN, SYT1, MYT1L, RIMBP2, ZMAT4, RAB3C, SYNPR, DLX6-AS1 | Neuron |
| FOLH1, MAG, MOBP, CLDN11, PLP1, KLK6, CNTN2, TF, UGT8, ST18, ERMN, XYLT1, SH3TC2, CNDP1, TMEM64 | Oligodendrocyte |
| GALR1, HAS2, NFYA, RGS13, TGFA, TNR, FPGT, MMRN1, CDH19, CA10, CRISPLD1 | Oligodendrocyte precursor cell |
